# Conditionally deleterious mutation load accumulates in genomic islands but can be purged with sufficient genotypic redundancy

**DOI:** 10.1101/2023.03.16.532996

**Authors:** Jonathan A. Mee, Bryce Carson, Sam Yeaman

**Author notes:** The authors wish to be identified to the reviewers.

## Abstract

Local adaptation frequently evolves in patches or environments that are connected via migration. In these cases, genomic regions that are linked to an adaptive locus experience reduced effective migration rates. Via individual-based simulations of a two-patch system, we show that this reduced effective migration results in the accumulation of conditionally deleterious mutations, but not universally deleterious mutations, adjacent to adaptive loci. When there is redundancy in the genetic basis of local adaptation (i.e. genotypic redundancy), turnover of adaptive loci allows conditionally deleterious mutation load to be purged. The amount of mutational load that accumulates adjacent to adaptive loci is dependent on redundancy, recombination rate, migration rate, population size, strength of selection, and the phenotypic effect size of adaptive alleles. Our results highlight the need to be cautious when interpreting patterns of local adaptation at the level of phenotype or fitness, as the genetic basis of local adaptation can be transient, and evolution may confer a degree of maladaptation to non-local environments.

## Introduction

Reciprocal transplant experiments commonly find evidence of local adaptation, where genotypes exhibit higher fitness when inhabiting their home environment and lower fitness when inhabiting other environments (Hedrick et al. 1976, Kawecki and Ebert 2004, Hereford 2009). Given its prevalence, it is important to understand how local adaptation evolves and whether it differs in any meaningful ways from global adaptation, where all populations of a species evolve towards the same optimum (Yeaman 2022). The fitness trade-offs that characterize local adaptation can arise due to three kinds of mutations (Figure 1; Fry 1996, Kawecki 1997, Anderson et al. 2013, Mee and Yeaman 2019): 1) antagonistic pleiotropy (AP), where alleles at a single locus are responsible for the trade-offs, with each allele being better adapted to one environment or the other; 2) conditionally beneficial (CB) mutations, where the derived allele has higher fitness than the wild-type in one environment and equal fitness in the other environment; 3) conditionally deleterious (CD) mutations, where the derived allele has lower fitness than the wild-type in one environment and equal fitness in the other environment. For the fitness trade-offs typical of local adaptation to arise via either CB or CD mutations, polymorphisms must be present at multiple loci, such that mutations with conditional effects conferring adaptation or non-local maladaptation in each environment are present at the same time. Hence, local adaptation can arise based on a single locus with an AP mutation, but the fitness trade-offs that characterize local adaptation can only arise via CB or CD mutations if multiple (i.e. at least two) loci are involved.

**Figure 1.**
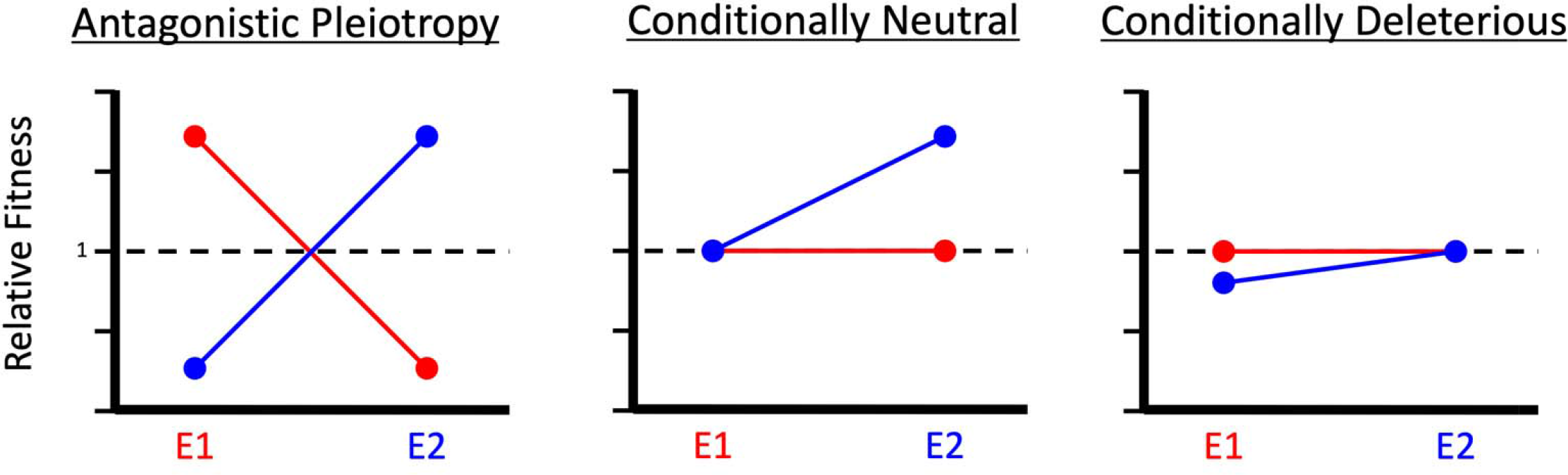
Genotype-by-environment interactions (i.e. fitness trade-offs) that characterize local adaptation can arise due to the three kinds of mutations depicted in this figure. Blue points and blue lines represent mutant alleles that contribute to local adaptation in environment 2 (E2) or non-local maladaptation in environment 1 (E1), whereas red points and red lines represent ancestral alleles that are adaptive or neutral in environment 1.

When there is antagonistic pleiotropy, both locally adapted alleles at a single locus will tend to be maintained at equilibrium provided migration is sufficiently restricted (Felsenstein 1976). By contrast, given sufficient time, CB mutations will tend to fix globally, while CD mutations will tend to be purged (Mee and Yeaman 2019). Previous analysis showed that a substantial load of CD mutations could be maintained under migration-selection balance, giving rise to fitness trade-offs typical of local adaptation (Mee and Yeaman 2019). While a substantial effect on fitness can arise from the transient segregation of many CD mutations, they typically don’t have strong differences in frequency between populations and therefore would not be expected to show strong genomic signatures of association (Mee and Yeaman 2019).

We explore the accumulation of conditionally deleterious (CD) mutations when they occur along with mutations that contribute to local adaptation via antagonistic pleiotropy. When the strength of selection on an AP locus is sufficient to overcome migration swamping, locally adapted alleles will tend to be maintained in environments where they are favoured (Lenormand and Raymond 2000). Substantial linkage disequilibrium can then emerge with other loci that are in tight physical linkage to the AP locus due to 1) reduced recombination between AP and CD mutations and 2) the persistent forcing effect of selection over migration. This results in a reduced effective migration rate or barrier effect at loci flanking the AP locus (Barton and Bengtsson 1986). Mee and Yeaman (2019) showed that the amount of load contributed by a CD locus would be approximately *L*_CD_ = *s*μ/*m*, where *s* is the strength of selection in the patch where it is disfavoured, *m* is the migration rate, and μ is the mutation rate. Given that an AP locus is expected to result in a reduced effective migration rate in its flanking regions, we would therefore expect that any CD loci within these regions could contribute higher load, as *L*_CD_ is higher with reduced *m*. Thus, it is possible that substantial CD load could arise near causal locally beneficial AP mutations driving, or contributing substantially to, the signatures of association we see in genome scans of local adaptation.

Linkage between AP loci and CD loci likely also has important implications related to constraints on adaptation. Segregating deleterious alleles can interfere with adaptation, which has been shown to result in slowed adaptation (Di et al. 2021). The effect of this interference may be particularly interesting in cases where there is genotypic redundancy in the genetic basis of local adaptation (*sensu* Láruson et al. 2020) such that fixation (or near-fixation) of adaptive alleles at only a subset of potentially adaptive loci is sufficient for local adaptation, but there are low-frequency segregating variants at multiple additional potentially adaptive loci. For example, if local adaptation is currently achieved via fixation (or near fixation) of an adaptive allele at one locus, but local adaptation could be achieved via fixation of an allele at any one of *n* other loci, then the current local adaptation represents one of *n* redundant possibilities. With this kind of redundancy, adaptive alleles may be transient, allowing for the purging of CD load when an adaptive allele rises in frequency at a different locus.

In this study, we explored two hypotheses related to the interaction of AP and CD loci. First, we hypothesized that substantial CD load will arise near AP mutations. Second, we hypothesized that redundancy in the genetic basis of local adaptation will have two effects: 1) transience of the genetic basis of local adaptation, and 2) purging of CD load. To explore these hypotheses, we used SLiM (Haller and Messer 2019) to simulate a two-patch model with either 1) a single AP locus on one chromosome to examine the accumulation of CD load, or 2) ten AP loci (one on each of ten chromosomes) to explore the effects of segregating redundancy (*sensu* Láruson et al. 2020) on the transience of the genetic basis of local adaptation and purging of CD load.

## Methods

Two versions of SLiM 3 (Haller and Messer 2019) were used to simulate the evolution of individuals distributed in two patches: v.3.3.2 and v.3.4.0. The version was dependent on when the simulations were conducted, and which contributor ran the simulations (details are provided in the README.md file in the DRYAD data repository associated with this study). We conducted two types of simulations. In the first type, which was intended primarily to address our hypothesis that substantial CD load will arise near AP mutations, each individual had a genome consisting of a single 100kb chromosome with 100 genes and a single antagonistic pleiotropy (AP) locus located in the middle of the 50^th^ gene (at locus 49,548); this type of simulation was ran with SLiM v.3.4.0 (Haller and Messer 2019). In the second type of simulation, which was intended primarily to address our hypothesis regarding the effects of genetic redundancy, each individual had a genome consisting of ten 100kb chromosomes with 100 genes and a single antagonistic pleiotropy (AP) locus located in the middle of the 50^th^ gene on four or more chromosomes; this type of simulation was ran with two SLiM versions, v.3.3.2 and v.3.4.0. The earlier version of SLiM 3 was used for simulations described in Carson (2022); the later version was used for additional simulations to study redundancy and allelic turnover (details in the DRYAD data repository). In the second type of simulation, there were up to 10 adaptive loci, and either the allelic effect size (zAP) or the number of adaptive loci (numAP) were important parameters affecting the degree of redundancy in the genetic basis of adaptation (see below). In both types of simulation (i.e. one chromosome or ten chromosomes), the per-basepair recombination rate, *R*, was set to 10^−7^, and the between-gene recombination rate, *r*, was set to either 10^−6^ (for a compact genome) or 10^−3^ (for a non-compact genome). The effect of the adaptive alleles on fitness (*W*) was determined with the following equation:

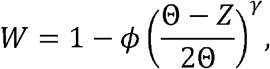

where *ϕ* is the strength of phenotypic selection (e.g. the fitness deficit in patch 1 of an individual perfectly adapted to patch 2), *Θ* is the phenotypic optimum in each patch (−1 in patch 1; 1 in patch 2), *Z* is the phenotypic value for a given individual, and *γ* is a shape parameter (which we set to 2 in all our simulations, resulting in a parabolic fitness function as shown in Figure 2).

**Figure 2.**
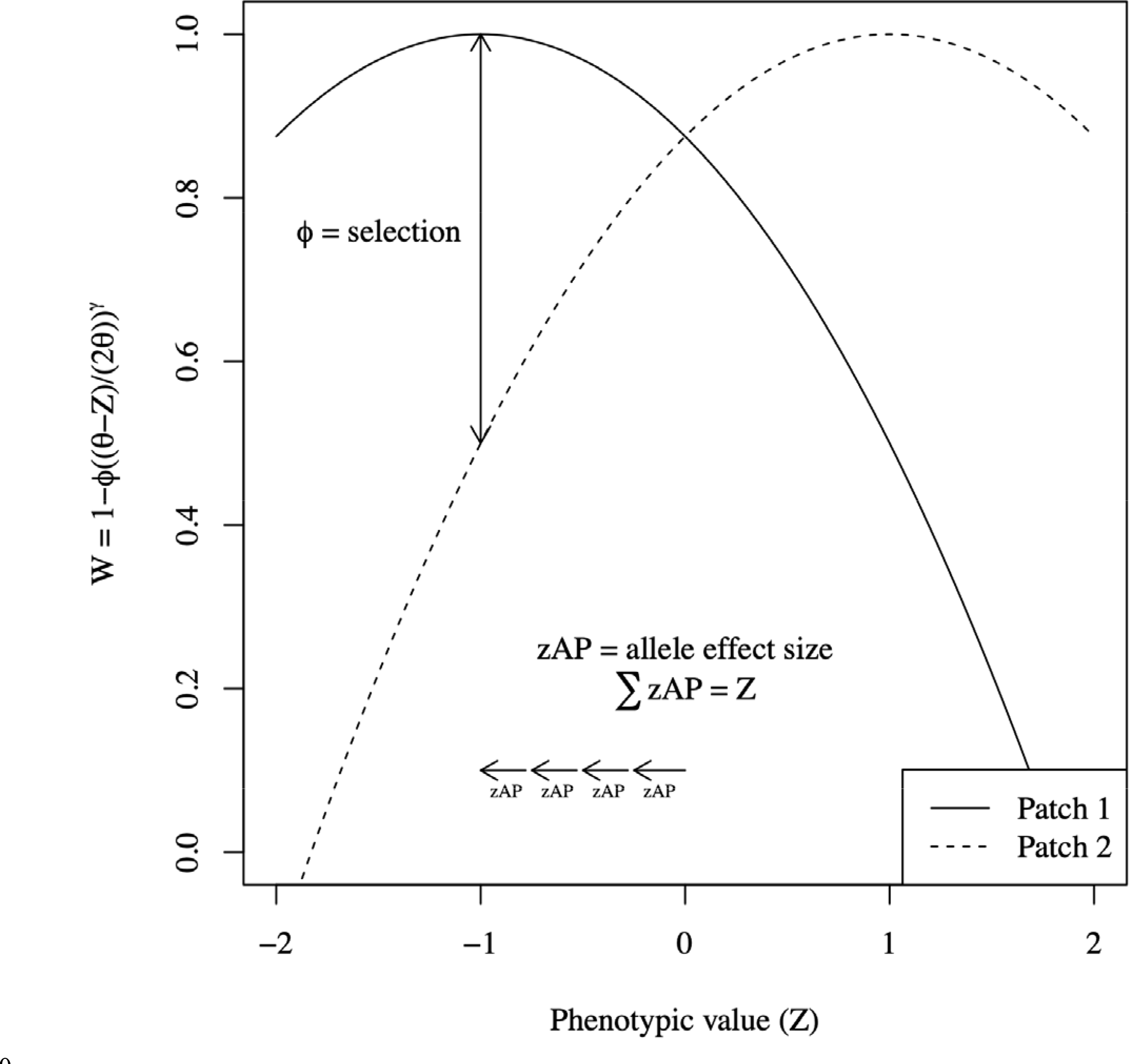
Fitness (W) is affected by an individual’s phenotype (Z) depending on the patch in which the individual resides. In our model, phenotype is affected by the additive effects of adaptive alleles (zAP). The phenotypic optimum (Θ) is set to -1 in patch 1, and 1 in patch 2. The figure depicts an individual with four alleles (at two or more loci) with effect size of -0.25, which is perfectly adapted to patch 1 (Z = -1) and suffers a fitness deficit of ϕ = 0.5 if it migrates to patch 2. The shape parameter, γ, is set to 2, as is the case for all our simulations.

Phenotypic effects of AP alleles (zAP) were additive across all loci. In the first type of simulation, with only a single AP locus, zAP was set to ±0.5 for all simulations (such that a -0.5 homozygote would be perfectly adapted in patch one, and a +0.5 homozygote would be perfectly adapted in patch 2). In the second type of simulation, with up to ten AP loci, zAP was set to one of the following values in each simulation: ±0.05, 0.0555, 0.0625, 0.0714, 0.0833, 0.1, 0.125, 0.1666, 0.25, or 0.5. In every simulation, there were three alleles that could arise by mutation at every AP locus: -zAP, 0, and +zAP. To ensure that the dynamics in our simulations were not limited by the availability of AP mutations, we set a (likely unrealistically) high AP mutation rate of μAP = 10^−4^ per locus, per generation. We made this decision because we wanted to focus on the accumulation of CD load *given* the existence of adaptive mutations. We are not drawing any conclusions about the pace of adaptation. Nonetheless, we did run simulations with lower AP mutation rates (μAP = 10^−5^, 10^−7^, 10^−8^, and 10^−9^) to explore the influence of AP mutation rate on turnover of adaptive loci. We set the strength of phenotypic selection to *ϕ* = 0.1 or *ϕ* = 0.5 for each simulation. Fitness effects of CD mutations were multiplicative. The effect size of each CD mutation was drawn from an exponential distribution, with a mean value of sCD specified for each parameter set depending on the between-patch migration rate (*m*) for that simulation, such that sCD = -*m*. The combined effect of CD and AP mutations on fitness was multiplicative, such that, for example, the fitness cost suffered by a patch-1-adapted migrant in patch 2 was multiplied against the fitness deficit resulting from that migrant’s load of CD mutations. The total calculation of relative fitness, following the algorithm SLiM defines and the fitness specification in the ten-chromosome simulations, is given in the following equation for a patch-1-adapted migrant individual in patch 2. *s* indicates the selection coefficient of a conditionally deleterious mutation or the phenotypic effect size of a mutation exhibiting antagonistic pleiotropy.

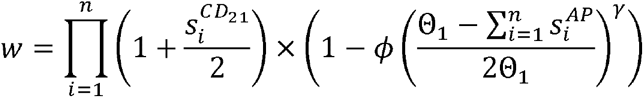

For all simulations, the CD mutation rate, μCD, was set to 10^−8^ per locus, per generation. All alleles (AP and CD) were codominant (h = 0.5). For comparison, we also included single-chromosome simulations with no AP mutations, with universally deleterious (as opposed to conditionally deleterious) mutations, and with no deleterious mutations at all.

At the beginning of every simulation run, for a 100,000 generation burn-in period, the between-patch migration rate was set to 0.5. Subsequently, the between-patch migration rate was set to *m* = 0.01 or 0.001. The number of individuals per patch was set to *N* = 1000 or 10,000. We ran each simulation for a total of 350,000 generations (including the burn-in), and recorded results every 5000 generations starting at the end of the burn-in period. We conducted ten replicate simulations for each parameter set.

## Results

### One chromosome, one adaptive locus

Substantial conditionally deleterious load accumulated adjacent to adaptive loci. We observed clear peaks in CD load in regions linked to the AP locus (Figure 3). In addition, the accumulation of the CD load was more tightly associated with the AP locus when migration rate was higher and when recombination rate was higher (Figure 3, Figure 4). The overall amount of deleterious load accumulated to a greater extent when phenotypic selection was stronger (Figure 4). Accumulation of deleterious load in the region around the AP locus only occurred with conditionally deleterious mutations, and not with universally deleterious mutations (Figure 3). The accumulation of the deleterious load did not occur to the same extent, nor in the same localized manner, when there were no AP mutations (Figure 3). Hence, the accumulation of deleterious load adjacent to an adaptive locus is a signature specific to conditionally deleterious mutations and is the result of linkage between adaptive alleles and conditionally deleterious alleles.

**Figure 3.**
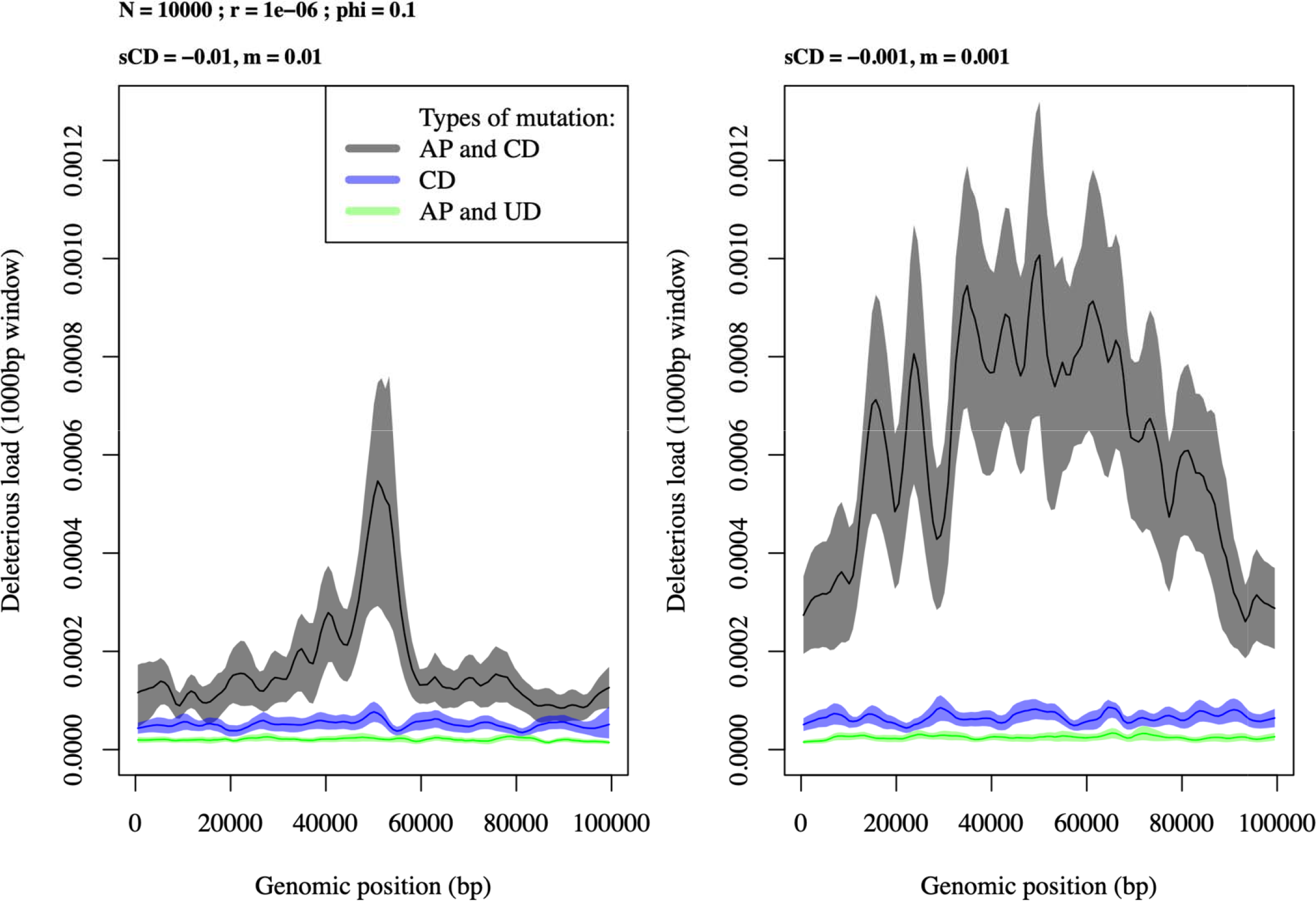
Deleterious load accumulates when the mutations are conditionally deleterious (CD), but not when mutations are universally deleterious (UD). In addition, conditionally deleterious load accumulates adjacent to the adaptive locus (the AP locus), which is in the center of each plot (at position 49,548 on the chromosome), and this CD load does not accumulate when there is no AP locus. Dark lines indicate the mean load across replicate simulations in a 1000bp sliding window, and the corresponding shading shows one standard error.

**Figure 4.**
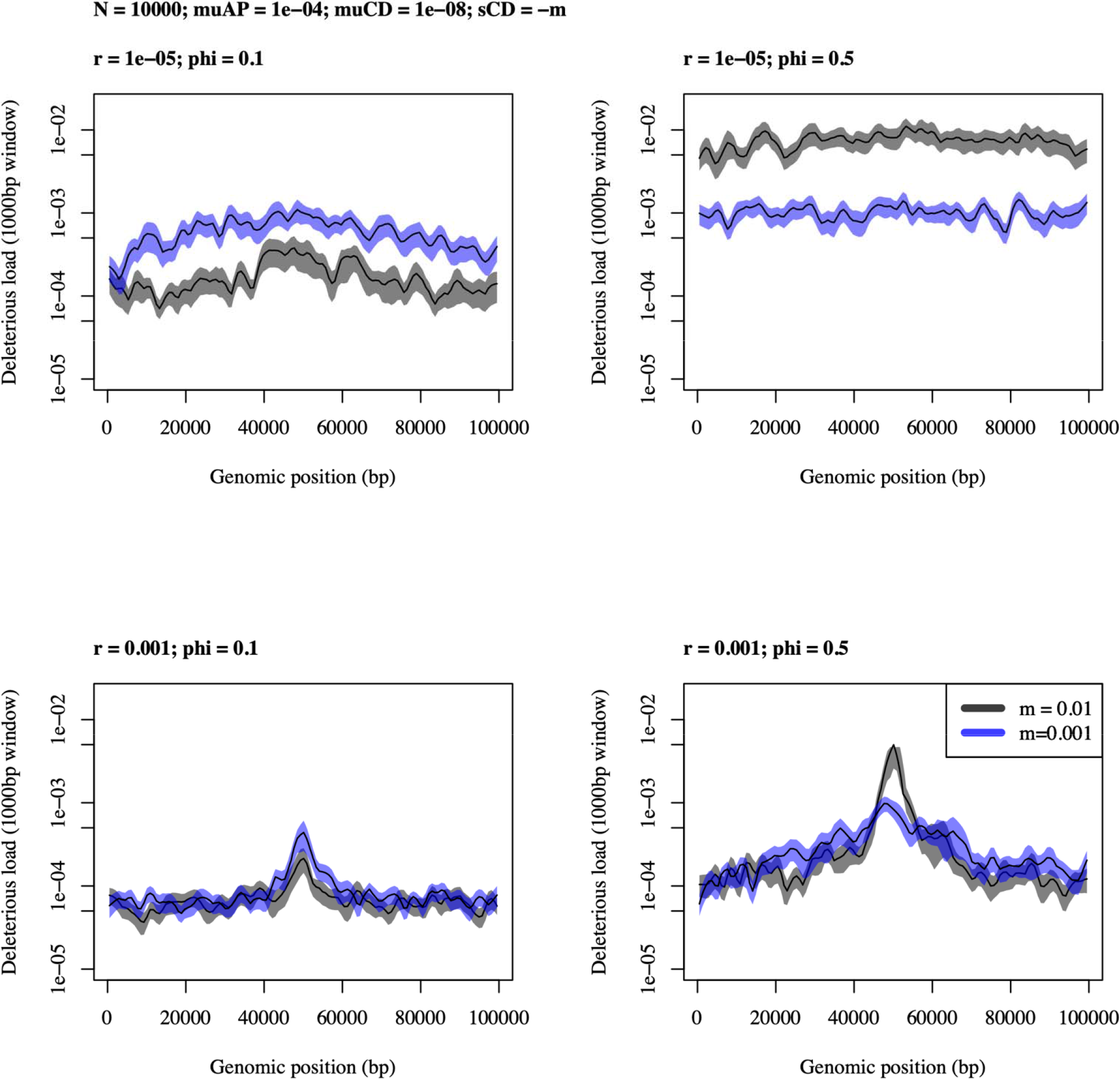
When genes are less densely packed along the chromosome (i.e. when the between-gene recombination rate, r, is high; top panels, r = 0.00001), the deleterious load is more easily purged across most of the chromosome, but accumulates strongly at loci linked to the adaptive locus (bottom panels, r = 0.001). Stronger selection on the adaptive phenotype (i.e. higher values of phi, right-hand panels) causes more accumulation of load, especially when migration rates are high. Dark blue or black lines indicate the mean load across replicate simulations in a 1000bp sliding window, and the corresponding shading shows one standard error.

The effect of migration rate on the accumulation of deleterious load interacted with the strength of the selection and with recombination rate (Figure 4). When selection was weak (*ϕ* = 0.1), higher migration rate resulted in reduced accumulation of load. When selection was strong (*ϕ* = 0.5), higher migration resulted in increased accumulation of load. When recombination was high (*r* = 10^−3^), the effect of selection on accumulated load (described in the previous two sentences) was confined to the region linked to the AP locus, whereas when recombination was low (*r* = 10^−5^), the effect of selection on accumulated load was observed across the chromosome.

### Ten chromosomes, ten adaptive loci

As in the simulations with one chromosome and one AP locus, deleterious load accumulated in regions linked to the AP loci when there were four to ten AP loci. The pattern of accumulated deleterious load adjacent to AP loci was clearest in the case with no redundancy, where adaptive allele effect size (zAP) was ±0.05 and there were ten adaptive loci, which necessitated adaptive alleles at all ten AP loci to reach the phenotypic optimum in a patch (Figure S1). In this case, as in the simulations with a single adaptive locus, higher migration rate reduced the accumulation of load when selection was weak. Also, there was less localization of the deleterious load, and overall lower deleterious load, when there were no AP mutations (Figure S1). Hence, the simluations with ten adaptive loci and no redundancy behaved similarly to the simulations with one chromosome and one adaptive locus.

We observed turnover in the adaptive loci when there was redundancy, so we explored whether changes in parameters that affect this turnover would change the maintenance of CD load, and whether the accumulation of CD load would in turn affect turnover. Genotypic redundancy can be altered by changes in the number of loci or the allele effect size (Yeaman 2015). We quantified turnover in each of our simulations by first calculating the standard deviation in the frequency of the adaptive allele on each chromosome across all sampled generations, then taking the average of these ten standard deviation values. We note that, because of the unrealistically high AP mutation rate in most of our simulations, we are not drawing conclusions about how much turnover in adaptive alleles we expect to see on a given time scale. We did, however, explore lower (and likely more realistic) AP mutations rates and found that turnover occurs even at low AP mutation rates (Figure S2). The amount of turnover was influenced by a combination of mutation rate, migration rate, allele effect size (zAP), and redundancy, but was not strongly affected by the presence/absence of CD mutations (Figure 5, Figure S2). This suggests that accumulation of CD load is not a strong driver of turnover of adaptive alleles.

**Figure 5.**
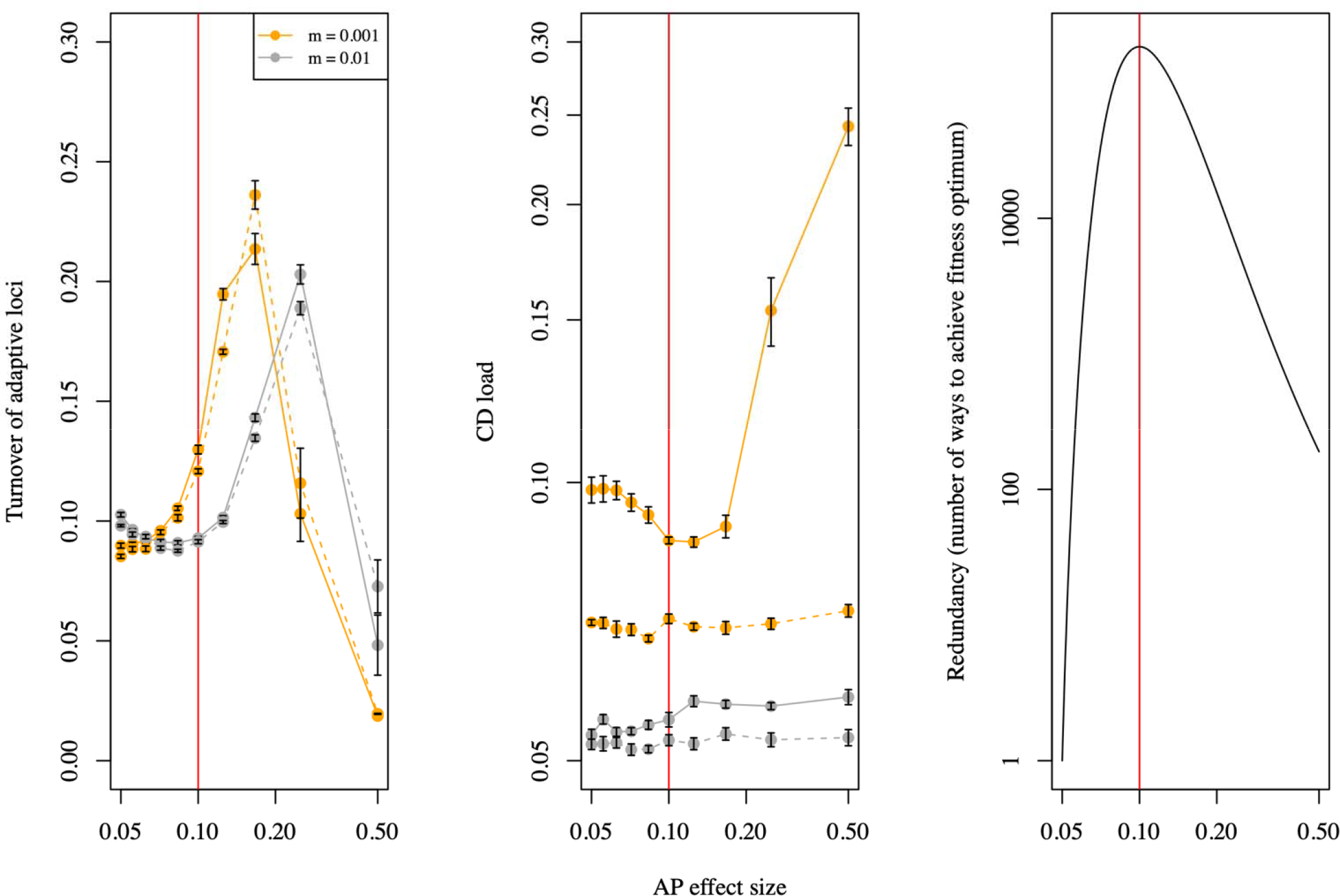
Turnover of adaptive loci (left panel) is the standard deviation, across all sampled generations, in the frequency of the adaptive alleles on each chromosome averaged across all chromosomes. CD load (middle panel) is the multiplicative load across all CD mutations in the genome (note log scale on both axes). Points show mean values across replicate simulations, and error bars show one standard error. Solid lines denote low between-gene recombination rate (r = 10^−6^), and dashed lines indicate high between-gene recombination rate (r = 10^−3^). The amount of genetic redundancy depends on the phenotypic effect size of the adaptive mutations (right panel). To be perfectly locally-adapted, the phenotypic effects across all alleles (AP effect size) must add up to 1 across all ten AP loci.

We explored two ways that changes in our parameters could alter redundancy, thereby affecting turnover and the accumulation of CD load: changing the allele effect size (zAP) and changing the number of loci. Because increasing the allele effect size changes the maintenance of local adaptation at the AP loci (Yeaman and Otto 2011), this resulted in complicated effects on turnover and CD load. Patterns were clearest at the low migration rate (*m* = 0.001), where conditionally deleterious load was purged when turnover of adaptive alleles was high, which resulted in a pattern of low overall CD load with high redundancy (Figure 5). In this case, because the number of loci is held constant, highest redundancy is achieved at intermediate allele effect sizes (Figure 5). At higher migration rates (*m* = 0.01), the relationship between turnover and purging of CD load is somewhat obscured by other effects associated with increasing allele effect sizes: stronger selection associated with larger alleles can entrain more of the chromosome in LD, which could increase the potential to maintain load. Thus, we see less clear relationships between redundancy, turnover, and the maintenance of CD load at higher migration rates. The pattern of reduced load with high genotypic redundancy was also evident when we held the effect size of adaptive mutations constant and varied the number of AP loci (Figure 6).

**Figure 6.**
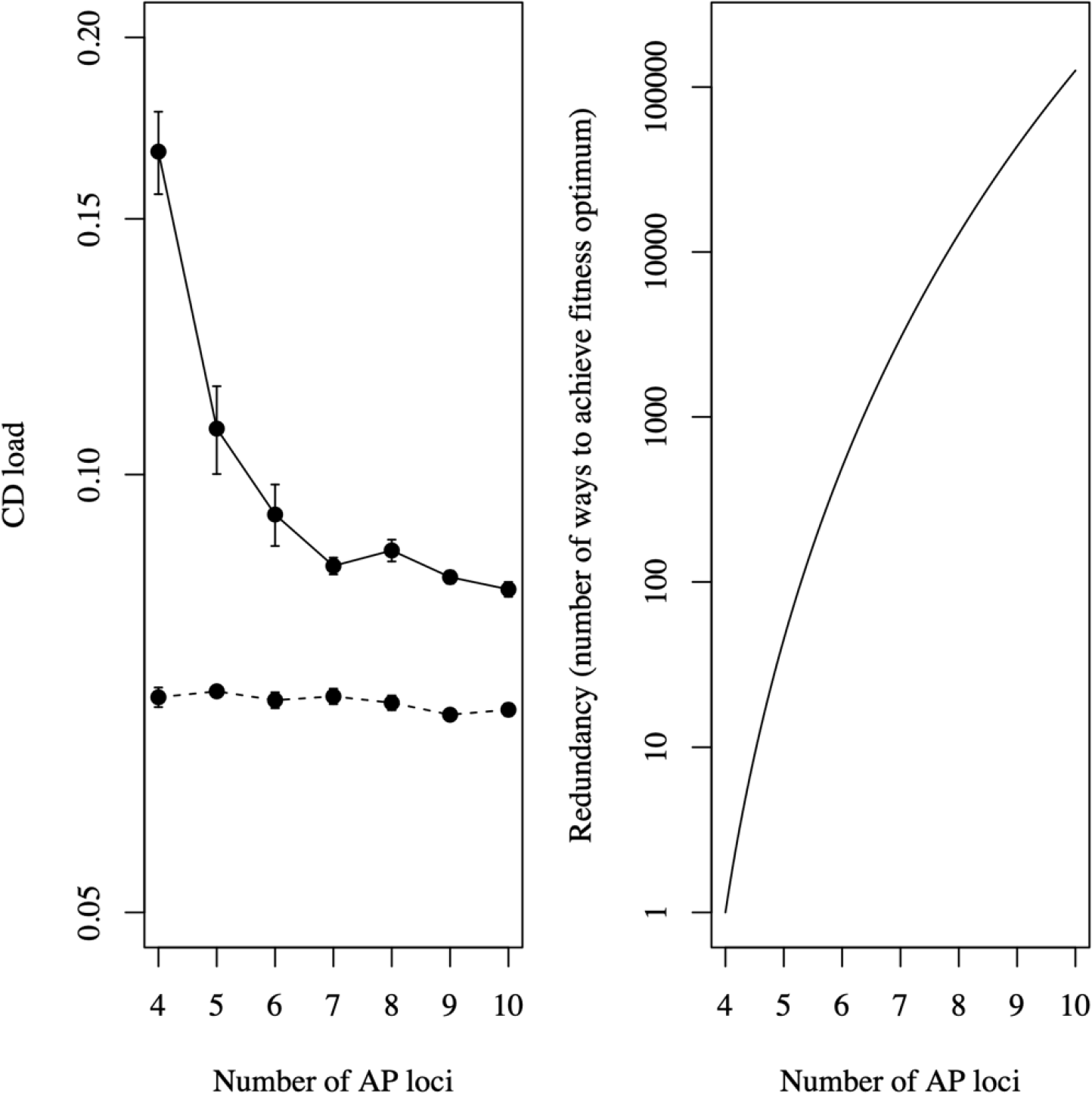
Conditionally deleterious load (left panel) is the multiplicative load across all CD mutations in the genome (note log scale on the y-axis). Points show mean values across replicate simulations, and error bars show one standard error. Solid lines denote low between-gene recombination rate (r = 10^−6^), and dashed lines indicate high between-gene recombination rate (r = 10^−3^). In this case, the effect size of AP mutations is held constant at zAP = +/- 0.125, and the amount of genetic redundancy depends on the number of AP loci (right panel). To be perfectly locally-adapted, an individual must carry eight adaptive alleles (e.g. must be homozygous for the adaptive allele at four loci). Other parameters are as follows: *N* = 10,000, *m* = 0.001, and *ϕ* = 0.1.

## Discussion

When a given locus drives local adaptation with gene flow, we have demonstrated that linkage with conditionally deleterious (CD) mutations causes the accumulation of deleterious load in its flanking regions. Importantly, the accumulation of linked deleterious load is specific to CD mutations as opposed to universally deleterious (UD) mutations. Unfortunately, empirical tests rarely have sufficient statistical power to identify whether a mutation is CD or UD because it is difficult to prove a negative: failing to detect a deleterious effect in all environments could simply be a type II error or a limitation of experimental design. Nonetheless, there is no reason to expect the accumulation of UD mutations around a locally adapted locus at equilibrium, so the observation of deleterious load concentrated adjacent to a locus involved in local adaptation would likely signal the involvement of CD mutations. One other plausible explanation for a localized enrichment of deleterious mutations is through hitchhiking during the initial spread of a new locally adapted mutation (Hartfield and Otto 2011), however such UD mutations would rapidly be purged once the mutation had established. Thus, it is possible to identify CD mutations via observations of linked deleterious load flanking an established locally adapted allele, and it would likely be fruitful to investigate well-characterized cases where local adaptation has evolved in the presence of gene flow. For example, *EPAS1* is strongly implicated in adaptation to high altitude in Tibetan populations (Zhang et al. 2021). Deleterious load linked to *EPAS1* could be identified via predictive software such as PROVEAN (Choi and Chan 2015, Sandell and Sharp 2022) or via a database of known non-synonymous SNPs in the human genome (e.g. Liu et al. 2011). Other cases where the genetic basis of local adaptation is well-characterized, and investigations of linked deleterious load may prove fruitful in the identification of CD mutations, include *Ectodysplasin* (*Eda*) in threespine stickleback (Schluter et al. 2021), Flowering time (*FT*) in *Arabidopsis thaliana, Helianthus annuus*, and many other plants (Izawa 2007; Auge et al. 2019; Todesco et al. 2020), and *Melanocortin-1 receptor* (*Mc1r*) in vertebrates (Manceau et al. 2010; Rosenblum et al. 2014).

A particularly good case for testing this theory is to look for enrichment in inversions involved in local adaptation, as they would be expected to accumulate CD mutations over a wider range of physical distance where recombination is reduced. Interestingly, a recent study in sunflower found that regions of the genome harbouring inversions tended to be enriched for non-synonymous mutations only in populations where both inversion haplotypes were segregating, but that no enrichment was found where the inversion haplotype was monomorphic (Huang et al. 2022). Given that they found no evidence of inversion overdominance (which could occur due to recessive UD mutations; Berdan et al. 2021) and inversion haplotypes instead tended to be found as homozygotes above the expected frequency, these observations are consistent with local adaptation driving maintenance of the inversions, leading to transient maintenance of CD load in the populations where migration-selection balance was operating.

Via our simulations that allowed the possibility of genotypic redundancy, we showed that 1) redundancy leads to turnover in the genetic basis of local adaptation, and that 2) turnover causes purging of the deleterious load that accumulates adjacent to adaptive loci. If, in a two-patch system, there are individuals in one patch whose local adaptation is based on a locus with linked CD load (locus A) and there are individuals in the same patch whose local adaptation is based on a locus without linked CD load (locus B), one might predict that occasional migration out of the patch might be sufficient to induce selection against the individuals with linked CD load at locus A, thereby favouring individuals with an adaptive allele at locus B and promoting turnover. Our results suggest, however, that the effect of CD mutations on the rate of turnover is small or negligible (Figure S2) and that the rate of turnover is determined primarily by genotypic redundancy and the effects of mutation rate, allele effect size, recombination rate, and migration rate on the adaptive locus itself. Understanding the contexts in which deleterious mutations are expected to drive patterns of turnover and genetic parallelism is an ongoing goal in the study of evolutionary genomics. For example, the accumulation of deleterious load has been hypothesized to drive sex chromosome turnovers (Blaser et al. 2014). Nonetheless, our results suggest that turnover is likely a feature of the genetic basis of local adaptation whenever there is genotypic redundancy, and the consequence of this turnover is the purging of CD load. This result should motivate empirical research on the extent of genotypic redundancy in cases of local adaptation, which will have implications on how often, and in what contexts, to expect the accumulation of CD load.

The existence of CD load linked to adaptive loci has implications for some approaches to the study of the genetic basis of local adaptation. The most obvious implication is in the identification of the locus (or loci) responsible for local adaptation based on mapping home-versus-away fitness differences. Any signal emerging from such a study may be caused both an adaptive effect (i.e. via a mutation at an AP locus) and a maladaptive effect (i.e. via mutations at linked CD loci), but these effects would likely be very difficult to disentangle. In studies that attempt to verify the genetic basis of local adaptation by using CRISPR to introduce a suspected adaptive allele into a naive genetic background, an otherwise-unexplained attenuated fitness effect of the CRISPR-inserted allele may be observed if the original background included linked CD load.

Our simulations highlight the complex and, in many cases, non-intuitive outcomes of interactions between gene flow, genetic drift, recombination rates, mutation rates, mutation effects, and natural selection. Our results add to the reasons to remain cautious in the interpretation of patterns of phenotypic or fitness differences among populations or demes, and to avoid simple explanations based on the primacy of any one mechanism (such as natural selection). The genetic basis of local adaptation can be transient, and adaptive evolution may confer a previously un(der)estimated degree of maladaptation to non-local environments.

## Acknowledgements

We thank Clément Rougeux for conversations and assistance with code development in early stages of this project. We thank the Digital Research Alliance of Canada for providing computing resources and support for many of the simulations described in this study. BC was supported by an NSERC USRA during the completion of this project. JAM used funds from an NSERC Discovery Grant to support and disseminate this project. SY had support from the Digital Research Alliance of Canada, NSERC Discovery, and Alberta Innovates.

## Data and Code Archiving

Single-chromosome simulation data was archived on DRYAD, with the generating SLiM scripts and supplemental code (R, Python, and BASH) on Zenodo. A large batch of ten-chromosome simulation output is hosted on the Federated Research Data Repository (FRDR), with the generating SLiM script, R scripts, and facilitating BASH code hosted on GitHub and archived on Zenodo. The remainder of the ten-chromosome data was archived on DRYAD, with the generating SLiM scripts and supplemental code (R, Python, and BASH) on Zenodo. FRDR archive: https://www.frdr-dfdr.ca/repo/dataset/88dae8ea-eb2f-43c2-8c3c-38ba5ca67fd1 DRYAD archive DOI: doi:10.5061/dryad.sf7m0cgb6 Private URL for DRYAD archive during peer review (Clicking the link immediately launches a download of the data files): https://datadryad.org/stash/share/sHeNumGe06liJerQH1Jo_jVjH2O44G-XUWgtAassdxU

---

**Figure S1.**
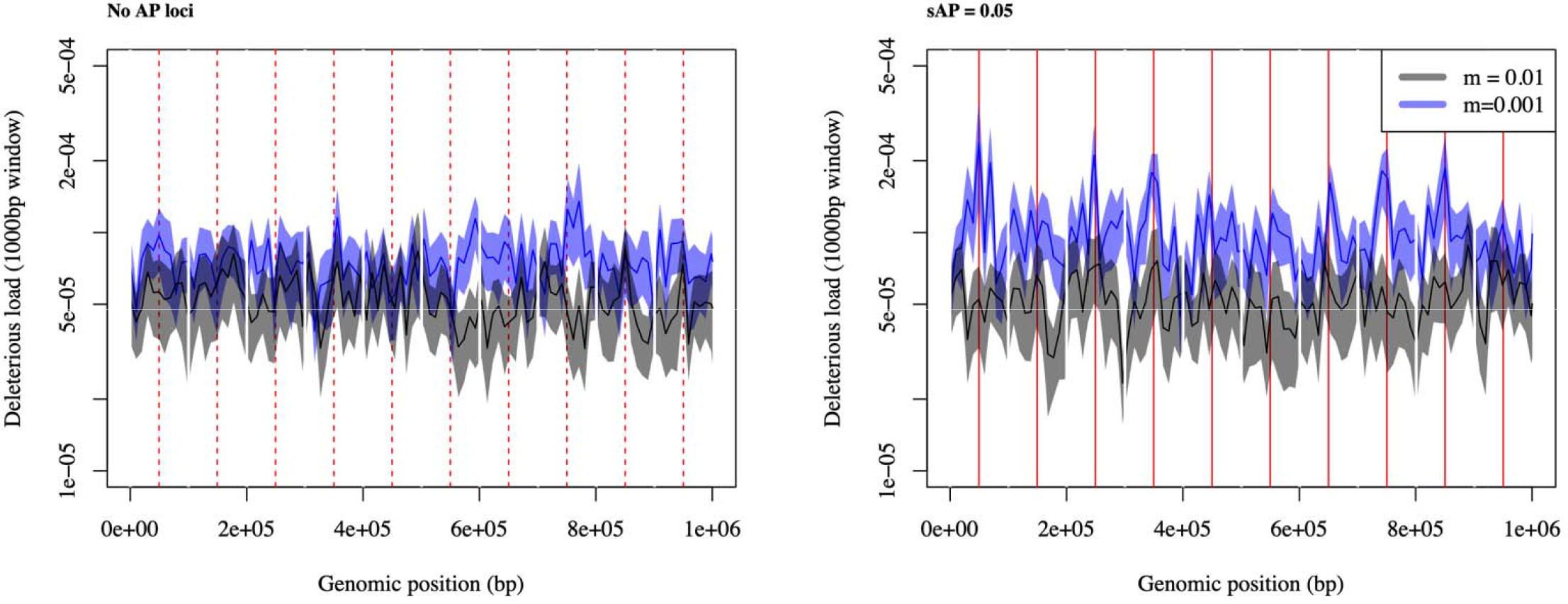
Load tends to be at its highest adjacent to the adaptive locus on every chromosome when every chromosome carries a single adaptive locus (located near the mid-point of each chromosome, indicated by red vertical lines), and when the effect size of adaptive alleles allows no redundancy such that every adaptive locus must carry an adaptive allele to be perfectly adapted (i.e. zAP = 0.05; right-hand panel). When there are no adaptive alleles (left-hand panel), load accumulates across the genome, and there is no tendency for load to be at its highest near the mid-points of the chromosomes (chromosome mid-points are indicated by vertical dashed lines). Dark blue or black lines indicate the mean load across replicate simulations in a 1000bp sliding window, and the corresponding shading shows one standard error.

**Figure S2.**
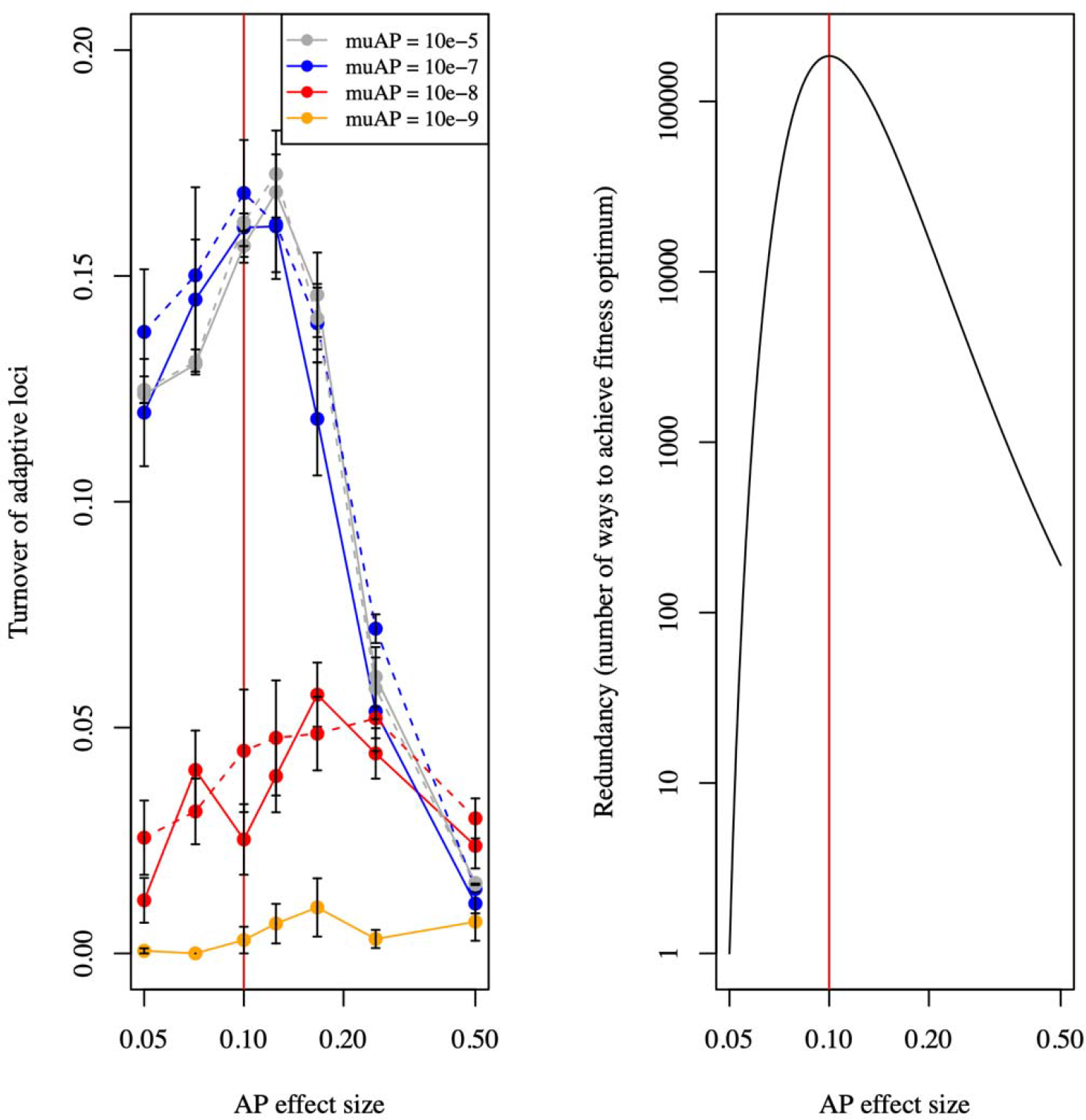
Turnover of adaptive loci (left panel) is the standard deviation, across all sampled generations, in the frequency of the adaptive alleles on each chromosome averaged across all chromosomes. Points show mean values across replicate simulations, and error bars show one standard error. Solid lines denote simulations without CD mutations, and dashed lines denote simulations with CD mutations. The amount of genetic redundancy depends on the phenotypic effect size of the adaptive mutations (right panel). To be perfectly locally-adapted, the phenotypic effects across all alleles (AP effect size) must add up to 1 across all ten AP loci.

